# Evaluation of redundancy analysis to identify signatures of local adaptation

**DOI:** 10.1101/258988

**Authors:** Thibaut Capblancq, Keurcien Luu, Michael G.B. Blum, Eric Bazin

**Author notes:** Deceased (Eric Bazin). Corresponding author: Thibaut Capblancq, 2233 Rue de la Piscine, 38041 Grenoble Cedex, France.

## Abstract

Ordination is a common tool in ecology that aims at representing complex biological information in a reduced space. In landscape genetics, ordination methods such as principal component analysis (PCA) have been used to detect adaptive variation based on genomic data. Taking advantage of environmental data in addition to genotype data, redundancy analysis (RDA) is another ordination approach that is useful to detect adaptive variation. This paper aims at proposing a test statistic based on RDA to search for loci under selection. We compare redundancy analysis to pcadapt, which is a nonconstrained ordination method, and to a latent factor mixed model (LFMM), which is a univariate genotype-environment association method. Individual-based simulations identify evolutionary scenarios where RDA genome scans have a greater statistical power than genome scans based on PCA. By constraining the analysis with environmental variables, RDA performs better than PCA in identifying adaptive variation when selection gradients are weakly correlated with population structure. Additionally, we show that if RDA and LFMM have a similar power to identify genetic markers associated with environmental variables, the RDA-based procedure has the advantage to identify the main selective gradients as a combination of environmental variables. To give a concrete illustration of RDA in population genomics, we apply this method to the detection of outliers and selective gradients on an SNP data set of *Populus trichocarpa* (Geraldes *et al.*, 2013). The RDA-based approach identifies the main selective gradient contrasting southern and coastal populations to northern and continental populations in the northwestern American coast.

## INTRODUCTION

Natural selection results from environmental pressures. The environment acts simultaneously on several characteristics, and these adaptive characteristics can be determined by a large number of alleles of small effects (Pritchard and Di Rienzo, 2010). Patterns of adaptive variation within species are usually studied through physiological, morphometric and fitness comparisons in common garden experiments. However, this is a difficult and constraining task, which can even be unrealistic to achieve in species with a long generation time. Recently, the development of next-generation sequencing (NGS) technologies has opened the possibility to access to a large amount of genetic variation across the genome. Therefore, relationships between genetic polymorphism, phenotypic variation and environmental variables can now be studied and quantified by in situ approaches (Steane *et al.*, 2014, Vangestel *et al.*, 2018).

By looking for loci that are either excessively differentiated between populations or significantly associated with environmental gradients, adaptive variation can be detected (Foll and Gaggiotti 2008, Frichot *et al.*, 2013, Vatsiou *et al.*, 2015, Duforet-Frebourg *et al.*, 2016, Hoban *et al.*, 2016).

In many genome scan procedures, outlier loci are identified based on genetic differentiation, assuming that adaptive alleles have increased genetic differentiation. A common statistic to evaluate genetic differentiation is F_st_ (Lewontin and Krakauer, 1973) and many likelihood or Bayesian methods use F_st_-related statistics to scan genomes for local adaptation (Bazin *et al.*, 2010, De Villemereuil and Gaggiotti, 2015, Foll and Gaggiotti, 2008, Whitlock and Lotterhos, 2015). One limitation of F_st_-based approaches is that they require defining discrete populations, whereas many species are not structured in different populations but display continuous genetic gradients (Martins *et al.* 2016). In these cases, defining populations can be challenging and can directly interfere with the result of the analysis (Yang *et al.*, 2012).

A solution to avoid such an issue is to use individual-based multivariate methods such as PCA that ascertain population structure. Duforet-Frebourg and colleagues (2016) proposed a genome scan method based on PCA (pcadapt), where outlier loci are the ones excessively correlated with one or more ordination axes (Duforet-Frebourg *et al.*, 2016). Simulations show that pcadapt has a similar power to model-based methods under island models and perform better when the simulated demographic model drifts from the island model, for instance under hierarchical population structure or isolation by distance patterns (Luu *et al.*, 2017). Nevertheless, one conundrum of such approaches is the difficulty to interpret ordination axes in terms of ecological meanings. These are usually tight to geographical axes (latitudinal or longitudinal), but they are not necessarily linked to an environmental variable such as temperature, drought, diet habit, etc. When environmental information exists, it has to be used, a posteriori, as a means of interpretation, but it is not involved in the inference process.

Genotype-environment association (GEA) methods are based on a different principle than methods based on genetic differentiation; they assume that adaptive loci are significantly associated with environmental variation. These methods aim at detecting alleles that are associated with environmental variables (e.g., temperature and drought) with the idea that these alleles may confer a selective advantage in some environment (Coop *et al.*, 2010, Frichot *et al.*, 2013). Most of these methods only use one predictor variable at a time, which can be problematic when the main selective gradient is unknown or when we want to disentangle multifactorial associations between genetic and environmental variation.

Some authors have proposed to use a constrained ordination procedure such as RDA (redundancy analysis) or CAPs (canonical analysis of principal coordinates) to combine the advantages of both multivariate methods and genotype-environment association procedures (Lasky *et al.*, 2012, DeKort *et al.*, 2014; Steane *et al.*, 2014, Hecht *et al.*, 2015). This family of methods subsequently aligns adaptive variation with environmental data in order to be able to identify the environmental gradients that are the most correlated with adaptive variation. In *Arabidopsis thaliana*, loci involved in adaptation to climate have been found using RDA (Lasky *et al.*, 2012). Outliers were identified as SNPs with the greatest squared loadings along the first RDA axes (i.e., those in the 0.5% tail). RDA can also be used to derive an adaptive index that predicts the performance of individuals in different environmental conditions. Steane *et al.* (2014) applied a similar approach (using canonical analysis of principal coordinates) to *Eucalyptus tricarpa* populations and showed that both detection of adaptive variation with RDA and common garden experiments provided evidence for local adaptation. Finally, in two recent studies, Forester and colleagues (2015 and 2017) tested the power of an RDA-based method for detecting signatures of local adaptation in a heterogeneous landscape (Forester *et al*, 2015) and in comparison with other constrained ordination methods (Forester *et al*, 2017). Using simulations, they found that an RDA-based method has a superior combination of low false positive and high true positive rates when compared to generalized linear models (GLM) or latent factor mixed models (LFMM).

In this paper, we build on these previous applications of RDA to develop a statistical test for searching loci under selection. We use simulations and real data sets to shed light on the conditions within which RDA is the most efficient and to document the possibilities given by RDA when studying landscape genomics. We also compare results of a genome scan based on RDA with genome scans obtained with *pcadapt* and LFMM (Frichot *et al.* 2013, Luu *et al.*, 2017). This investigation gives the opportunity to show the advantages of a constrained ordination method such as RDA in comparison with nonconstrained ordination method (PCA) and univariate genotype-environment association method (LFMM). Finally, to illustrate the pertinence of such a method in a concrete example, we apply it to a *Populus trichocarpa* SNP dataset obtained from Geraldes *et al.* (2013). We found a large overlap between the outlier loci detected with RDA and the loci previously detected with *Fst* based methods (Geraldes *et al.*, 2013). In addition, RDA revealed what the main selective gradients are along the natural populations of *Populus trichocarpa* in the northwestern coast of North America.

## MATERIALS AND METHODS

### 1. Genome scan using an RDA-based approach

Redundancy analysis (RDA) was first introduced by Rao (1964). It is the extension of multiple regressions to the modeling of multivariate response data (Legendre and Legendre 2012, section 11.1). The data are separated in two sets, a response matrix Y of the variable to be explained (e.g., species abundance in a set of sites; m sites and n species) and an explanatory matrix X (e.g., a set of environmental variables within each site; m sites and p environment). In the following analysis, loci replace species abundances and individuals replace sites. RDA seeks to project the genetic variation between individuals that is explained by environmental data on a reduced space. RDA assumes that there are linear relationships between the response (Y) and explanatory (X) variables.

The analysis starts using a classical RDA procedure performed with the function *rda* of the R package *“vegan”* (Oksanen *et al.*, 2015). An individual genetic dataset is used as the response matrix Y, and a set of environmental variables is used as explanatory matrix X. We assume in the following that genotypes are coded using the values 0,1, or 2 that correspond to homozygote for the most frequent allele, heterozygote and homozygote for the less frequent allele. RDA amounts at constructing a matrix Y’ of fitted genetic values estimated from the regression of each locus by the environmental variables and at performing principal component analysis on the matrix Y’ (Legendre and Legendre 2012, section 11.1). After this constrained ordination step, we follow the methodology implemented in *pcadapt* to find outlier loci (Luu *et al.*, 2017). First, we recover the loci loadings from the RDA analysis. Only the loadings of the most informative ordination axis are kept for the rest of the procedure. The number of axes used (K) is determined by looking at the amount of information retained on the different axes of the RDA. A Mahalanobis distance D is then computed for each locus to identify loci showing extreme D values compared to the rest of the SNPs. A Mahalanobis distance is a multidimensional generalization of the idea of measuring how many standard deviations is a point from an average point. Computation of the Mahalanobis distance uses the *covRob* function of the “*robust*” R package (Wang *et al.*, 2014). Mahalanobis distances are distributed as chi-squared distribution with K degrees of freedom after correcting with the genomic inflation factor (Luu *et al.*, 2017). Inflation factors are constant values that are used to rescale chi-square statistics in order to limit inflation due to diverse confounding factors (François et al 2016). We then adjust the resulting p-values for the false discovery rate (FDR) by computing q-values with the “qvalue” R package (Dabney and Storey, 2011). A locus is considered as an outlier if its q-value is less than 10%, meaning that 10% or less of the identified outliers could be false positives. The complete procedure is described in Fig. 1 and our R script is available in the supplementary material (Script S1).

**Figure 1:**
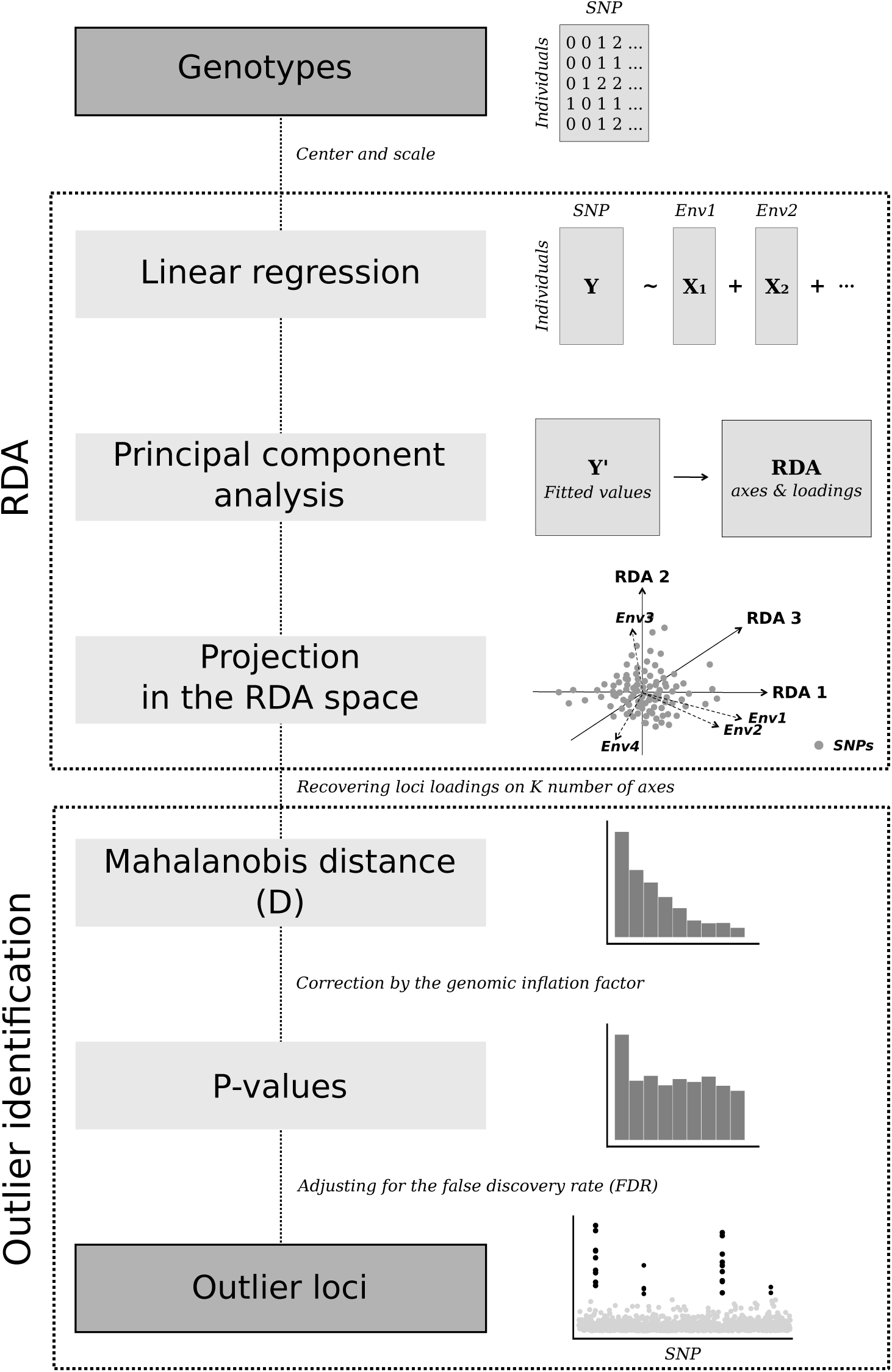
Schematic representation of the RDA-based procedure for searching loci under selection.

To evaluate the capacity of the RDA-based approach to detect loci underlying environmental selective pressure, we first test the method with simulated datasets. A total of 100 independent runs of simulations were used (see below for details). We evaluate the power of the procedure based on RDA to detect positive loci (loci simulated under selection) and to return a controlled number of false positives (neutral loci detected as outliers by the analysis). Then, to emphasize the utility of this method, we compared it to the PCA-based procedure implemented in *pcadapt* that does not use environmental variables (Luu *et al.*, 2017). In the same vein, we compare the multivariate RDA-based procedure to LFMM, which is an univariate genotype-environment association method (Frichot *et al.*, 2013). This method processes one environmental variable at a time, whereas we wanted to evaluate to what extent different techniques can disentangle covariation between multiple environmental gradients and multilocus genetic variation. To tackle this issue, we performed the LFMM analysis with a set of composite environmental variables corresponding to the first two PCs of a PCA performed on the environmental simulated variables (Frichot *et al.*, 2013). It gave us orthogonal composite environmental variables for which we tested covariation with genetic variability through the LFMM procedure. The LFMM analysis was launched with only one latent factor (K = 1), taking into account that simulations have been made without any population divergence.

### 2. Simulated data

We used simulations performed with the simuPop python library (Peng *et al.*, 2005). A lattice of 8x8 populations is simulated (i.e., 64 populations in total). Each population is initialized with 200 diploid individuals with random genotypes. Migration is set to 0.1, resulting in an isolation-by-distance pattern across the species range. Loci are assumed to be biallelic SNPs. The allele frequency of the whole population is initialized at 0.5. 1000 loci are simulated in total and they are separated in 200 chunks of 5 SNPs in physical linkage with the recombination rate between adjacent loci fixed at 0.1. Three different quantitative traits are coded by a group of 10 different loci (quantitative trait locus, QTL). The first quantitative trait is coded by the loci 1, 11, 21, …, 91. The trait value is simply the sum of the genotype values and therefore can take values between 0 and 20. We add to each trait a random noise (nonheritable variation) drawn from a normal distribution *N*(0,2). The second quantitative trait is coded by loci 101, 111, …, 191 and the third is coded by loci 201, 211, …, 291. Each quantitative trait is therefore coded by 10 independent SNPs (QTL), resulting in a total of 30 causal SNPs among the 1000. Selection can have an effect on linked loci, for instance, loci 2, 3, 4 and 5 can be impacted by selection on locus 1. However, recombination is high enough (0.1) to expect a limited linkage effect. We have defined 10 different environmental variables. The first one determines the selective pressure on trait 1, the second one on trait 2 and the third one on trait 3. The first environment variable is a quadratic gradient coded by function *env1* = −(cos(*θ*)*(i-3.5))^2^ −(sin *θ*)*(j-3.5))^2^ + 18, *θ* = *π*/2, with i and j being the population indicator on the 8x8 lattice. The second one is a linear plan gradient coded by function *env2* = h*cos(*θ*)*(i-1) + h*sin(*θ*)*(j-1) + *k* with *h* = 2,*θ* = *π*/4 and *k*=3. The third environment variable simulates a coarse environment with value *env*3 = 2 for all populations except population (i,j) = (2,2), (2,3), (3,2), (3,3), (6,2), (6,3), (7,2), (7,3), (2,6), (2,7), (3,6), (3,7), (6,6), (6,7), (7,6), (7,7), for which *env*3 = 18. Env4, env5 and env6 have exactly the same equation as env1, env2 and env3, respectively, except for a noise term explained below. The remaining 4 environment variables are similar to env2 but with different values of h and *θ*. Env7 has *h* = 2, *θ* = 0 and *k* = 3. Env8 has *h* = 2, *θ* = *π*/4 and *k* = 0. Env9 has *h* = 1, *θ* = *π*/4 and *k*=4. Env10 has *h*=0.5, *θ* = *π*/4 and *k* = 8. Graphical representation of the mean environmental value is given in Fig. 2. Environmental variables 4, 5 and 6 have, respectively, the same mean value spatial distribution as environmental variables 1, 2 and 3. The environmental equation gives a mean value of the environment variable. To avoid collinearity between environmental variables, we added noise by drawing an environment value within a normal distribution N(*μ* = env, *σ* = 1). The fitness for each trait is set to be –exp(x-env)^2/(2*ω^2), with *x* being the quantitative trait value, *env* the environmental value and *ω* is the defining selection strength and has been set to 20, which was found to be enough for loci to be often detected. To get the overall fitness for a given individual, the fitness associated with each trait is multiplied. Fitness values are used to determine the number of offspring during the simulations, which have been launched for 500 generations. At the end of the simulation, we sampled 10 individuals per population resulting in 640 individuals with 1000 SNP-like loci.

**Figure 2:**
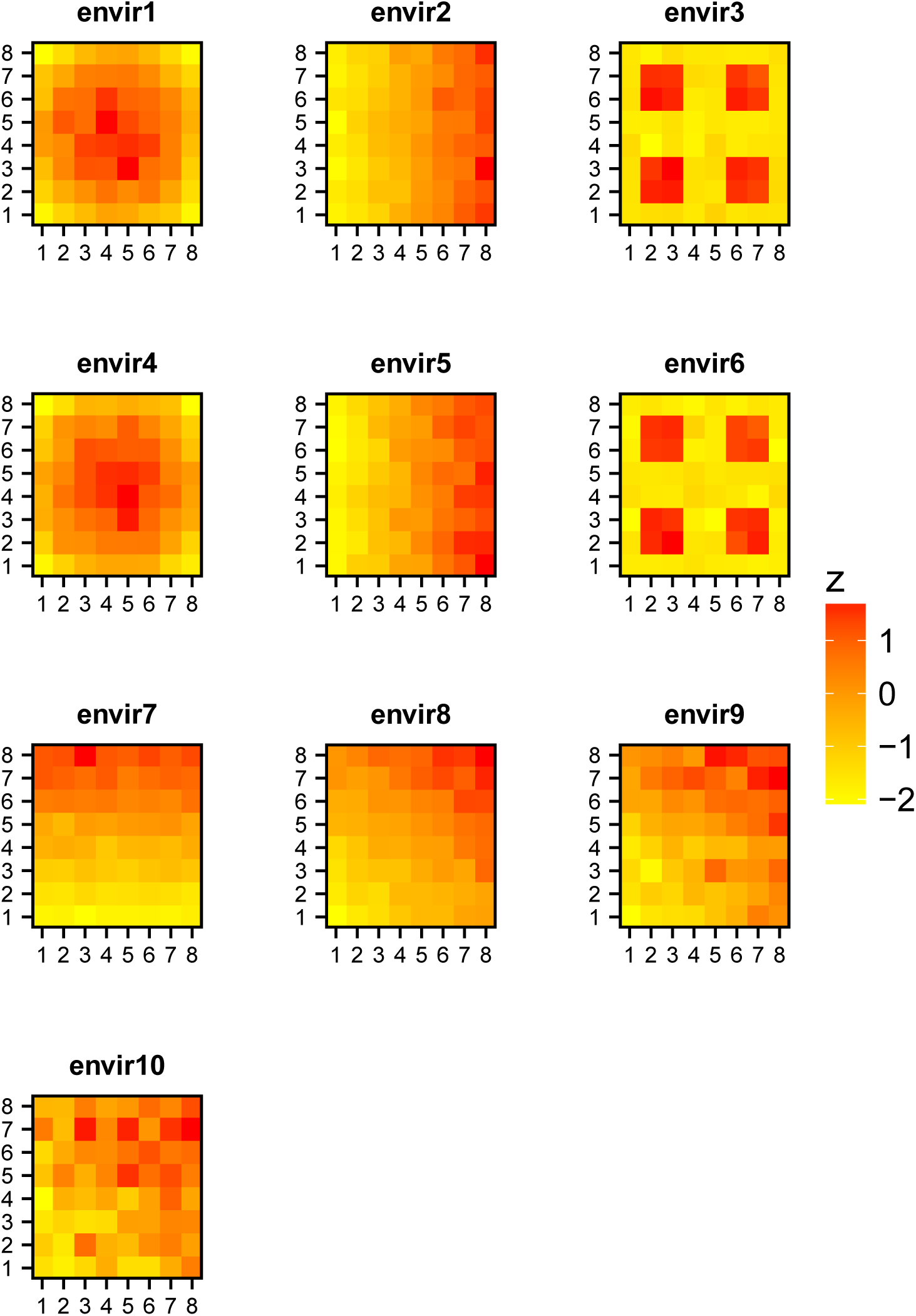
Graphical representations of mean environmental values for environments 1 to 10. Equation determining the environment is given in the main manuscript. Mean environmental values for environments 4, 5 and 6 are, respectively, equivalent to environments 1, 2 and 3.

### 3. Real dataset

The *Populus trichocarpa* dataset is a sample of 424 individuals genotyped on 33,070 SNPs from 25 drainages (i.e., topographic units separated by watershed barriers) (Geraldes *et al.*, 2014). Genotyping of each accession was performed with a 34K Populus SNP array targeting 34,131 SNPs mostly within (plus 2 kb upstream and downstream) 3543 genes (Geraldes *et al.*, 2013). A total of 21 climatic variables are available on each sampling site (Table S1). From the 33,070 SNPs, we removed the SNPs with missing values and a minor allele frequency below 5% resulting in 17,224 SNPs.

We explored the adaptive genetic variability of the *Populus trichocarpa* dataset using PCA-based (*pcadapt*) and RDA-based method using the 21 environmental variables as explanatory variables (Table S2). Similar to the simulations above, we created two environmental composite variables by keeping the two first axes of a PCA on the 21 environmental variables. We then used the latent factor mixed model (LFMM) procedure on these composite variables and with one latent factor (K=1). We finally compared the list of outliers found with these three approaches to the ones found using *Fst* based methods (Geraldes *et al.*, 2014).

We then performed a second RDA with only the loci previously found as outliers (q-values < 0.1). This set of outliers gives an “adaptively enriched genetic space” (Steane *et al.*, 2014) and using a second RDA on these specific loci, we can identify the environmental variables that are the most correlated with “putatively” adaptive variation. Using only outlier loci avoids potential interferences brought by the “neutral” genetic variation during the alignment between genetic and environmental variation performed by RDA. We then used the scores of the different *Populus trichocarpa* populations along the first two RDA axes to build composite indexes, which correspond to the main environmental pressures driving adaptive genetic variation. The individuals’ scores on the axes of the adaptively enriched genetic RDA can be interpreted as cursors of individual genetic adaptation to the environmental variables associated with these axes. We thus represented the mean RDA1 and RDA2 individual score for each of the 133 populations on a map. It allows visualizing the adaptive landscape across the sampling area.

## RESULTS

### 1. Genome scan on simulated dataset using RDA, pcadapt and LFMM methods

The procedure was first performed on one simulation (sim1) with pcadapt, RDA and LFMM approaches. To perform RDA, we considered environmental variables 1–10 as explanatory variables. For pcadapt and RDA, we retained the first four ordination axes to compute Mahalanobis distances as indicated by the scree plots (Fig. S1). In parallel, we independently launched LFMM analysis with the first two PCs resulting from a PCA applied to environmental variables (Fig. S2). We then estimate that a locus is considered as an outlier by LFMM if there is a significant correlation with at least one of the two composite variables.

The software package *pcadapt* is successful at detecting QTL2 SNPs (Fig. 3 topleft) but fails entirely at detecting QTL1 and QTL3 SNPs. The RDA-based approach also detects QTL2 SNPS and most of the QTL1 SNPs in contrast to *pcadapt* (Fig. 3 topright). The LFMM method is also able to detect most of the QTL1 and all the QTL2 SNPs. These SNPs are detected when respectively regressing the genetic dataset with the first and the second environmental composite variables (Fig. 3 bottom). For the SNPs associated with QTL3, the three methods have negligible power to detect them.

**Figure 3:**
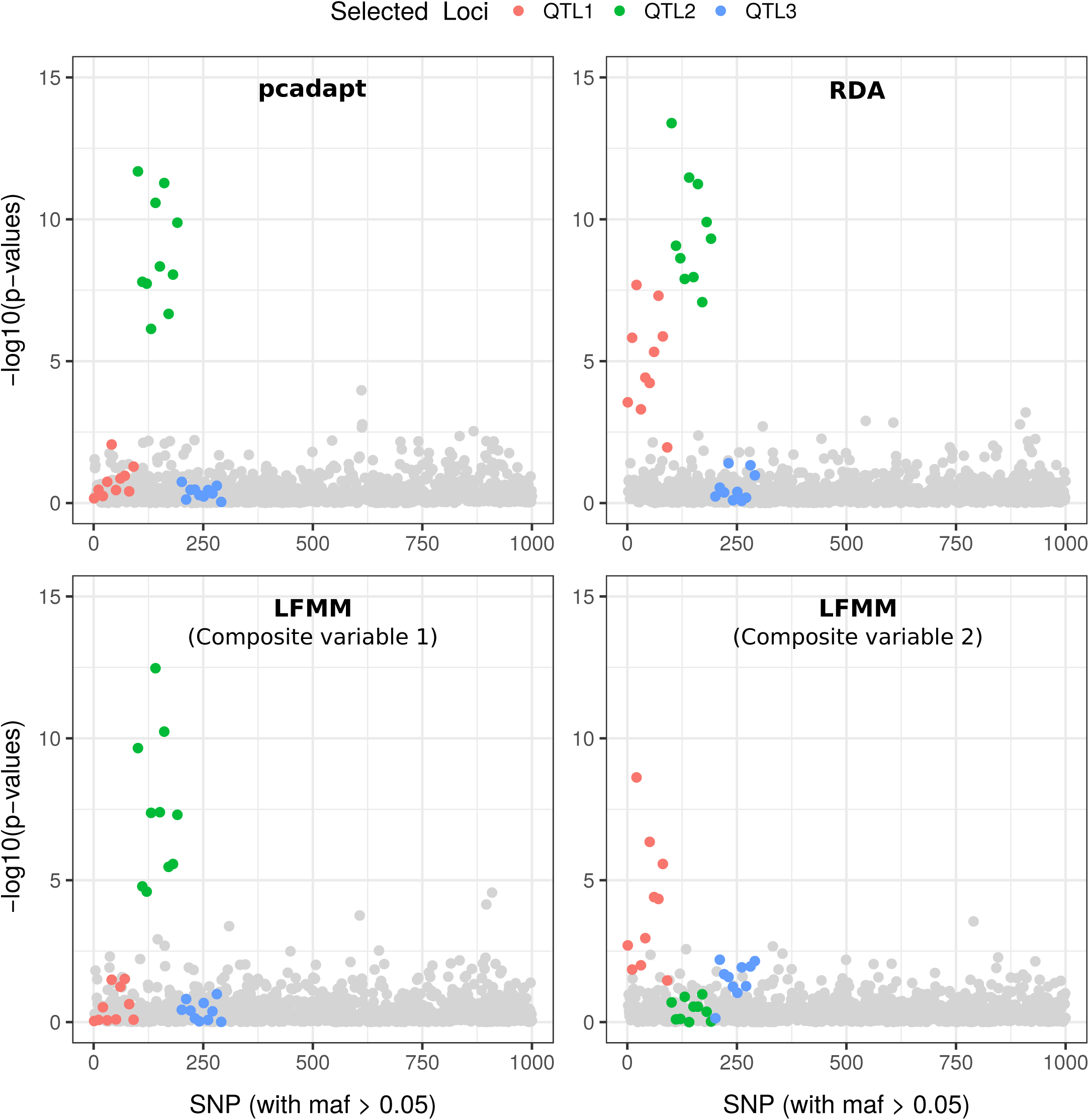
Manhattan plots of the results obtained with *pcadapt* (upper left panel), RDA (upper right panel), and LFMM (lower panels) for one simulated dataset.

When looking at the projection of each SNP in the RDA space, we can detect links between QTL SNPs and environmental variables (Fig. 4). RDA1 is strongly correlated to env2 and env5 and the QTL2 SNP show extreme scores on this axis compared to the other SNPs, especially the neutral ones (Fig. 4A). RDA2 is also associated with env7, env8, env9 and env10, which had not been used as direct selective drivers during the simulations. RDA3 is correlated to env1 and env4 corresponding to QTL1 SNPs and there is also a weak association between RDA4 (env3 and env6) and the SNPs associated with QTL3 (Fig. 4B & 3C).

**Figure 4:**
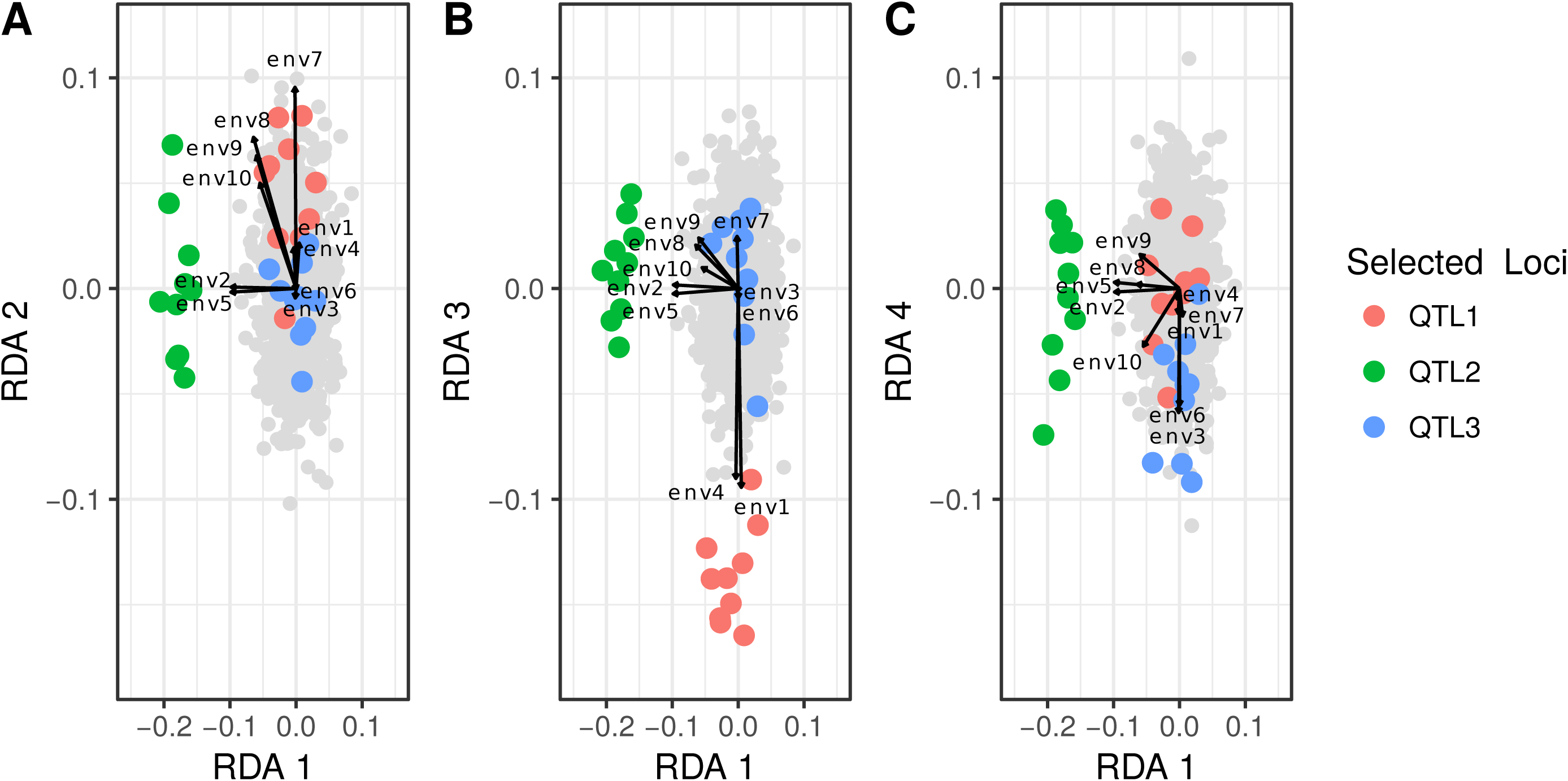
Projection of SNPs and environmental variables into the RDA space.

When looking at the percentage of environmental variance explained by the RDA and PCA axes, we find that the first four RDA axes explain more variance of each environmental variable than the first four axes of the PCA (Fig. 5). The difference is especially true for env1 and env4, which are strongly associated with the first four axes of the RDA, whereas PCA axes show an almost null correlation. The variables env7, env8, env9 and env10 are also correlated to the first four axes of the RDA, whereas PCs show far less correlation with these environments. Finally, RDA axes show more correlation with the env3 and env6 variables than PCs (R^2^ = ~0.30 vs R^2^ = ~0). Furthermore, each RDA axis, from the first one to the fourth one, is associated with more than one environmental variable, whereas only one PC (PC1) is associated with env2 and env5 (Fig. S3).

**Figure 5:**
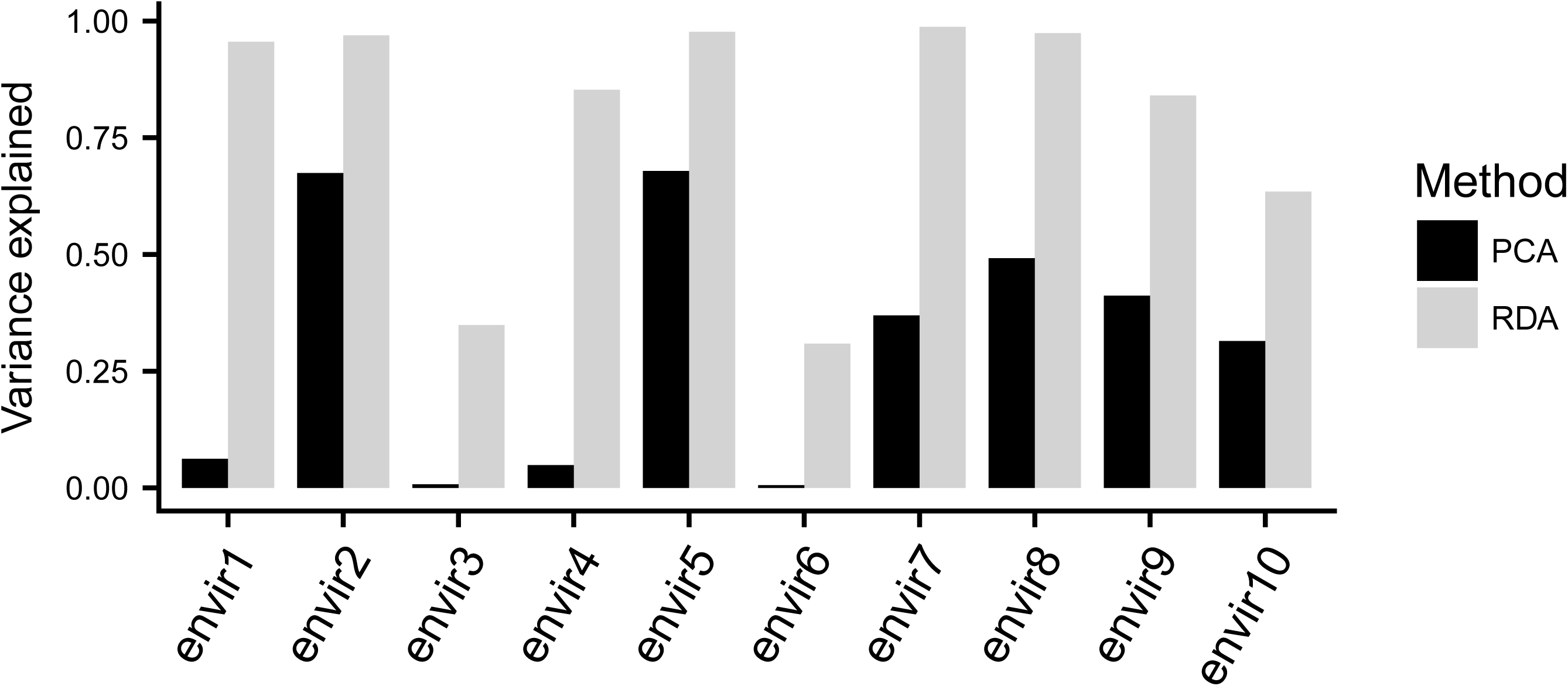
Proportion of environmental variance explained by the first four ordination axes using RDA or PCA. The values correspond to the r-squared of the regression between the environmental variable and the first four axes of the multivariate analysis.

Finally, over the 100 simulations, we measured the average FDR and power for *pcadapt*, LFMM and RDA (Fig. 6). The three methods correctly identify almost all of the QTL2 SNPs (Fig. 6A). RDA-based analysis and LFMM are more powerful than *pcadapt* to identify QTL1 SNPs with, respectively, 75%, 48% and less than 10% of true identification. For QTL3, *pcadapt* is not able to find any of the SNPs under selection, whereas RDA reaches 24% of true identification and LFMM reaches 22%. The three methods properly control for the false discovery rate because the proportion of false discoveries obtained with *pcadapt*, LFMM and the RDA-based methods are, respectively, 5%, 2.3% and 2.3% when controlling for a nominal error rate of 5%.

**Figure 6:**
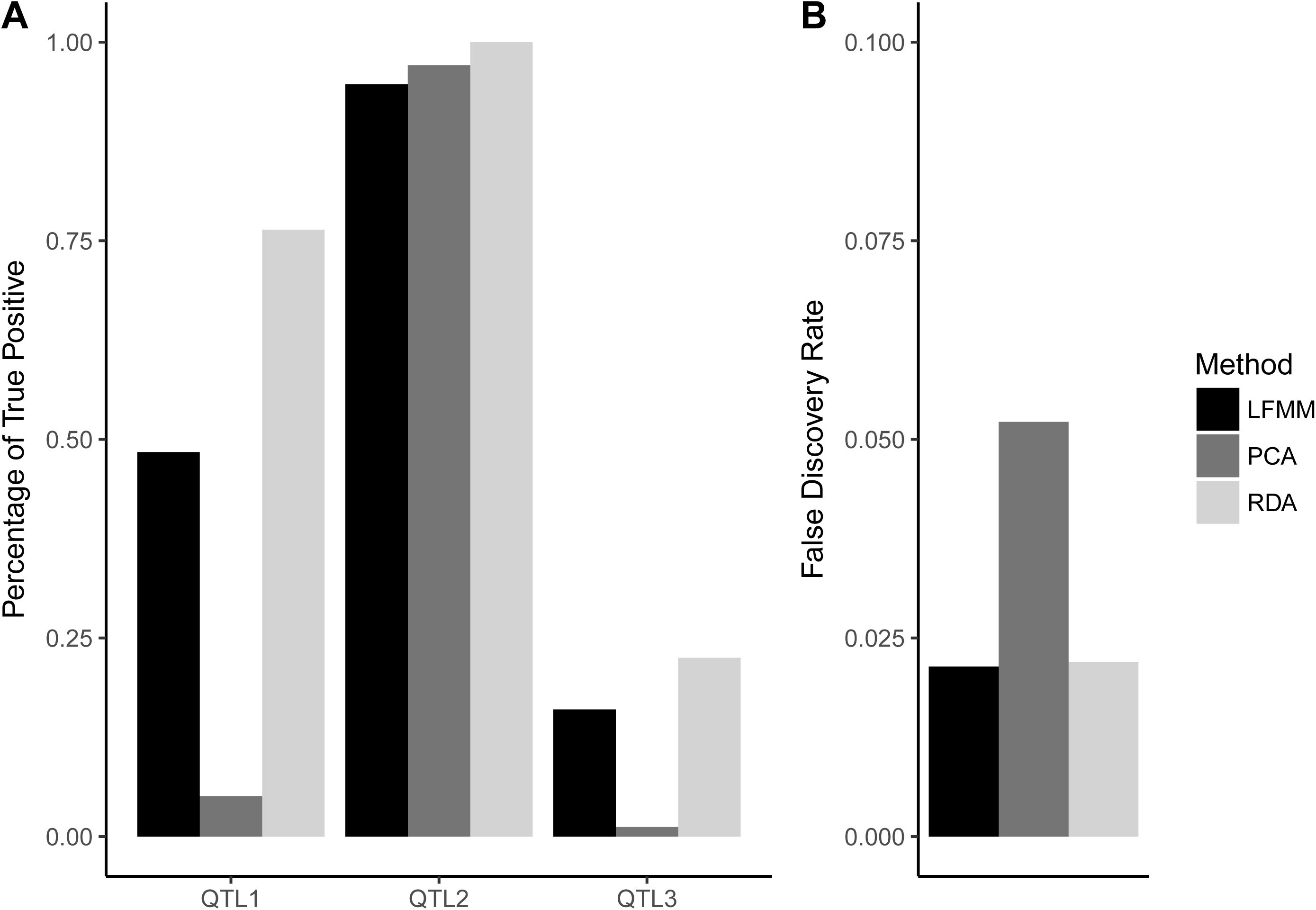
Statistical power obtained with RDA, *pcadapt*, and LFMM. Estimate of power corresponds to an average over 100 simulated datasets (error bars are displayed). Power (A) is given separately for loci coding for quantitative traits 1, 2 and 3. False Discovery Rate (FDR) is given for all methods (B).

### 2. Populus trichocarpa

For the RDA and *pcadapt* analysis on the *P. trichocarpa* dataset, we retained respectively *K*=3 and *K*=6 axes as indicated by the scree plots (Fig. S4) and we used the two first axes of a PCA on environmental variables as composite variables for the LFMM procedure (Fig. S5). Analysis with an FDR of 0.1 provides a list of 151 outliers for RDA, 117 for *pcadapt* and 36 for LFMM. Interestingly, the three methods have only 9 outliers in common and 117, 77 and 11 SNPs are outliers, respectively, specific to RDA, *pcadapt* and LFMM.

When we compared RDA genome scan results with the outliers found by Geraldes *et al.* (2014) based on Fdist, BayeScan and bayenv methods, we found a substantial overlap between them (Fig. 7). Among the 151 outliers found with the RDA-based method, 69 have already been picked up by *Fst*-based methods (see Table S3). Interestingly, the loci showing the lowest p-values with the RDA analysis are also pointed out as outliers by the *Fst* and *pcadapt* methods, but not by the LFMM procedure (Fig. 7). However, the RDA-based genome scan found 72 SNPs that no other methods, including *pcadapt*, detected as outliers.

**Figure 7:**
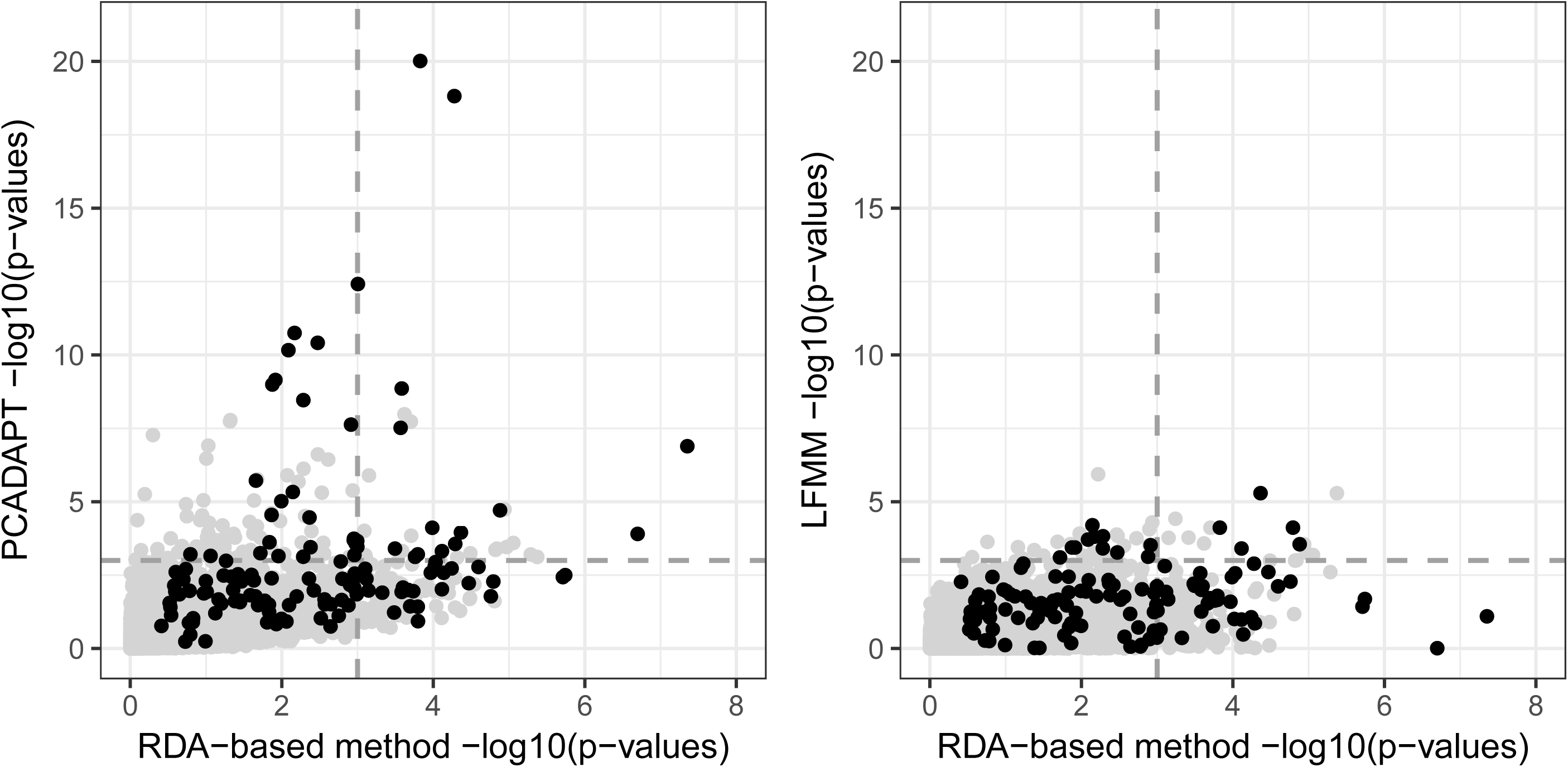
Scatter plot of the p-values returned by RDA and *pcadapt* procedures (A) and the p-values returned by RDA and LFMM procedure (B) for the *Populus trichocarpa* dataset. Black dots correspond to loci identified as outliers by *Fst* methods in Geraldes *et al.* (2014). The dashed lines indicate a 10–3 p-value with RDA and *pcadapt* or LFMM.

When we performed a new RDA on the set of 151 outlier loci as detected by the RDA-based genome scan method (adaptive enriched genetic space), we found that the two first axes keep most (respectively 40% and 18%) of the adaptive genetic variance among the populations (Fig. 8A). These two axes correspond to the two principal selective pressures in the sampling area. RDA1 is correlated with several environmental variables used in the analysis (Fig. 8A), including DD_0, DD18, MAT, MCMT, and EMT, which correspond to temperature indexes. RDA1 is also correlated to moisture variables (e.g., AHM or MAP), continentality (TD) and flowering period length (bFFP and eFFP). This axis summarizes a gradient of global climatic variation encountered in the sampling area with southern and coastal populations opposed to northern and continental populations (Fig. 8B). On the other side, RDA2 also correlates with moisture and temperature variables such as AHM, MAP, TD, MCMT, and eFFP and contrasts individuals from coastal locations to individuals from inland areas (Fig. 8C).

**Figure 8:**
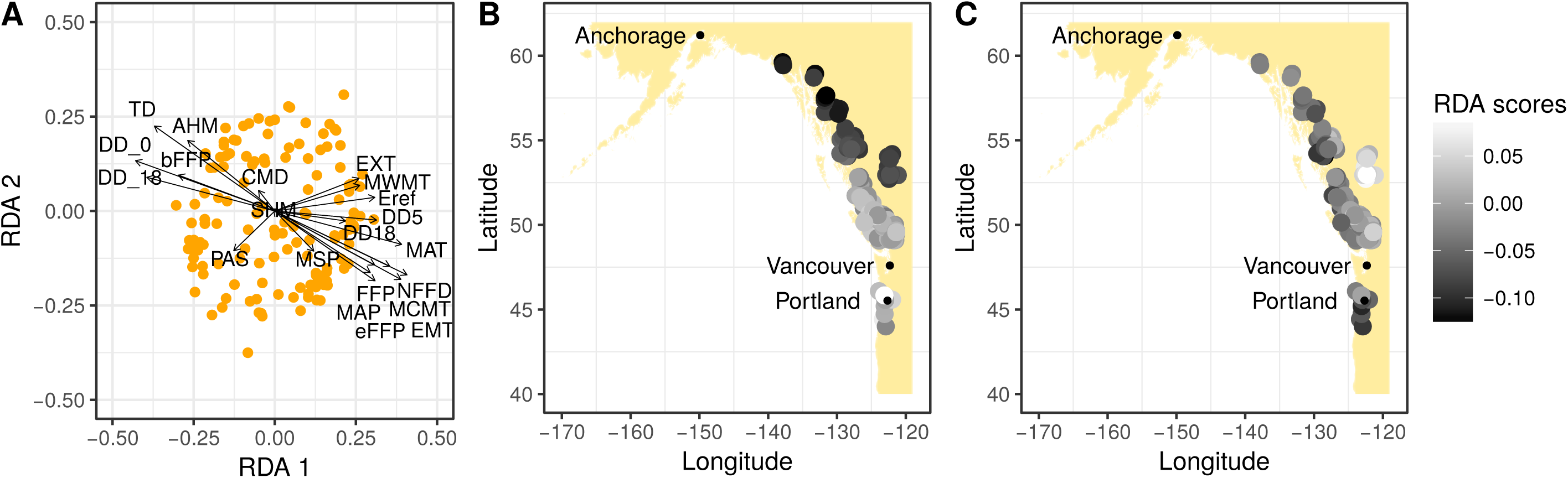
Projection of SNPs and environmental variables in the RDA performed on the adaptively enriched genetic space (151 loci) of *P. trichocarpa* dataset (A). The first two axes of the RDA projection represent 40% (RDA1) and 18% (RDA2) of the explained variance. (B) and (C) show respective spatial representations of RDA1 and RDA2 axes scores in the sampling area. The points represent the 133 sampled populations and their color depends on the mean of individual RDA1 (B) and RDA2 (C) scores in the populations.

## DISCUSSION

Genome scans based on genetic differentiation, as implemented in *pcadap*t, detect true positives when environmental gradients related to adaption are correlated to population structure. For instance, QTL2 SNPs are adaptive along an environmental gradient that is correlated with population structure (see Fig. 2) and they are detected by *pcadapt* (Fig. 6). By contrast, when the environmental gradient is not correlated to population structure (QTL3 and QTL1 to a lesser extent) it fails to detect adaptive SNPs. In this case, PCA ordination is not capable of orienting genetic variation into the direction of environment 1 or 3. Fig. 6 shows that RDA has a larger statistical power than *pcadapt* to detect QTL1 SNPs by taking advantage of information from environmental local conditions. It can be attributed to a better alignment between the genetic space and the environmental variable improving the power to detect true positives. This better alignment is easily visible when looking at the PCA and RDA correlation with environmental variables (Fig. 5 & Fig. S3). PCA is only capable of identifying axes, which correspond to maximum genetic differentiation (PC1 and PC2), whereas RDA is also able to capture the genetic variability associated with other environmental variables, such as env1 (RDA3) and to a lesser extent env3 (RDA4).

Genotype-environment approaches (GEA) explicitly take into account the relationship between genotypes and the environment (Joost *et al.* 2007, de Villemereuil *et al.* 2014, Lotterhos and Whitlock 2015). We showed that univariate GEA, implemented in LFMM, and multivariate GEA, based on RDA, can show similar results in identifying adaptive variation although RDA is more powerful than LFMM for QTL1 SNPs (Fig. 6). As suggested in the literature, we limited the number of environmental predictors during the LFMM procedure by using environmental PCA axes as composite variables (Frichot *et al.*, 2013). We then only used the two first PCs, but adding the third and fourth ones could potentially increase the capacity of LFMM to detect QTL1 and QTL3 SNPs. It points out o ne of the advantages of a multivariate procedure (e.g., RDA) compared to univariate GEA because multivariate methods take all environmental variation into account at the same time and can simultaneously detect associations between different sets of loci and different sets of environmental variables (Forester *et al.*, 2017).

Both RDA and LFMM fail to detect QTL3 outliers in, respectively, 75% and 80% of the simulations (Fig. 6), possibly because of the additional difficulty in detecting adaptive SNPs on a coarse environment compared to a smooth environmental gradient as in environment 1 and 2. LFMM and the RDA-based genome scan rely on linear association between the environment and allelic abundance (0, 1 or 2). Patchy selective pressure, such as env3, is difficult to identify with a correlative approach. Forester and colleagues (2015) already found that heterogeneous environmental selection decreases the capacity of RDA to identify the loci under selection. The strength of selection and the level of allelic dispersion during the simulations strongly influence the statistical power of RDA; weak selective strength and strong dispersion are the less favorable conditions (Forester *et al.*, 2015). Nonetheless, our simulations plead in favor of using a constrained ordination method when relevant environmental variables are available in order to both orientate the ordination axis in the direction of environmental gradients and to account for most of the environmental variation. This result confirms that constrained ordination shows a superior combination of low false positive and high true positives rates across all levels of selection (Forester *et al.*, 2017).

During our RDA-based procedure, we use Mahalanobis distances as a test statistic to identify genetic variation significantly associated with environmental gradients. This distance is computed based on the scores of more than one RDA axis and can thus condense the information of several RDA axes in one statistic. Some other studies have used different RDA-based statistics to identify the loci under putative selection. They often used a value estimated independently for the different RDA axes, such as the loadings of the loci along the axis (e.g., extreme squared loadings for Latsky *et al.*, 2012) and considering a threshold such as ± 3 SD (Forester *et al.*, 2017). Mahalanobis distances can capture complex selection patterns where allelic variation depends on more than one environmental gradient. This statistic is appropriate to identify selective pressures on natural populations, which usually encompass multifactorial gradients of selection (Salmela, 2014).

One application of ordination methods is to detect selective gradients. The RDA-based approach succeeded very well in decomposing the QTL SNPs-environments associations along the different axes, at least in our simulations. In our example, we succeeded in dissociating the outliers linked to the different environmental variables, and thus to precisely identify which set of variables drives each part of the adaptive genetic variability (Fig. 4). RDA1 is strongly associated with env2, RDA2 with env7 (collinear with isolation by distance pattern in the grid), RDA3 with env1 and RDA4 with env3. As expected, the correlated environments are also strongly associated with these respective axes. This is reflecting the fact that it is difficult to distinguish among several collinear variables which environmental variable has a causal effect on the individual fitness. However, it is often sufficient to identify the combination of environment variables having a strong association with adaptive variation without precisely knowing the underlying mechanical process. These simulations serve as a proof of concept to show that ordination axes can provide holistic measures of genomic adaptation (Steane *et al.*, 2014). Under this idea, scores of individuals on one RDA axis reflect their degree of adaptation along this axis. RDA produces useful visualization of gradients of genetic adaptation (Fig. 8).

From the analysis of *Populus trichocarpa*, RDA found numerous outlier loci (72) that are not detected by any other genome scan method, including *pcadapt* and LFMM (Fig. 7 and Table S3). These results support the simulation analysis conclusions that the RDA-based genome scan is able to pick up more genetic markers in relation to environmental gradients than *pcadapt*. It is, however, noticeable that a proportion of outliers are shared between PCA and RDA-based methods (29), highlighting the fact that both approaches are partially similar. The difficulty in interpreting the PCA axes is compensated by the fact that it is a blind method in regard to environmental data, so it can pick up genes that we will miss using RDA because some crucial environmental data are lacking. In the same vein, half of the outlier loci found with the LFMM procedure (14 over 33 loci) are common with RDA results (Fig. 7 and Table S3). The two methods rely on linear regression between environmental and genetic variability, and by using PCs as composite environmental variables in the LFMM procedure, we even add some similarity between the two approaches.

After focusing the RDA analysis on the loci showing significant association with environmental variables (151 outliers), we identified the predominant environmental gradients structuring the genetic adaptation of *Populus trichocarpa*. The main one is a composite climatic variation (RDA1 in Fig. 8), gathering precipitation, temperature and seasonality variations. It clearly contrasts the populations coming from the extreme northern part of the sampling area (Alaska) and the population coming from the inland locations (higher altitudes) to the populations close to the coast in British Columbia (Canada) and the southern populations, near Portland, Oregon (USA). This result is in accord with the conclusions of Geraldes *et al.* (2014). In their study, they point out the need of adaptation to seasonality, photoperiod and frost events for *Populus trichocarpa* along its continuous distribution from 44 to almost 60 degrees of latitude. RDA2 is associated with moisture regimes and points out a gradient of adaptation between coastal locations, strongly watered, and inland populations, receiving less precipitation (Fig. 8). These two environmental gradients are the strongest selective pressures driving *Populus trichocarpa* climatic adaptation in northwest America, considering the set of environmental variables available and the scale of the analysis. To go further, these composite indexes could also serve to predict (i) provenances that would perform well in common gardens; (ii) patterns of adaptation to local climate across the geographic range; or (iii) future adaptive landscapes in the context of climate change.

## ACKNOWLEDGEMENTS

We would like to dedicate this paper to the memory of our colleague and beloved friend Eric Bazin (1977–2017) who passed away too soon.

TC acknowledges support from the ANR project APPATS (ANR-15-CE02–0004) and MB acknowledges the Grenoble Alpes Data Institute, which is supported by the French National Research Agency under the “Investissements d’avenir” program (ANR-15-IDEX-02). MB also acknowledges support from the ANR project AGRHUM (ANR-14-CE02–0003–01).

## DATA ACCESSIBILITY STATEMENT

- Simulations datasets are available at the Dryad Digital Repository at the following address: https://datadryad.org/review?doi=doi:10.5061/dryad.1s7v5
- Scripts used for simulations and *Populus trichocarpa* analysis are available on Github: https://github.com/Capblancq/RDA-genome-scan
- *Populus trichocarpa* SNP data are available on Dryad: http://datadryad.org/resource/doi:10.5061/dryad.1051d

## AUTHOR CONTRIBUTIONS

E. Bazin designed the study. E.B. and T. Capblancq performed the analysis and treatments. M. Blum, E.B. and T.C. wrote the manuscript, and all authors contributed substantially to revisions.

## SUPPORTING INFORMATION

**Script S1:** Detailed script of the RDA-based genome scan analysis.

**Table S1:** Environmental variables used in the *Populus trichocarpa* analysis.

**Table S2:** Climatic variable values for the 133 *Populus trichocarpa* sampled populations.

**Table S3:** Comparison between RDA, *pcadapt*, *fdist* and *bayescan* outlier loci.

**Figure S1:** Scree plot for PCA and RDA analysis of the simulation 1 dataset.

**Figure S2:** PCA and scree plot of PCA on environmental variables of the first simulation.

**Figure S3:** Heatmap of the R^2^ resulting from linear regression between the four first axes of PCA and RDA analysis and the environmental variables.

**Figure S4:** Scree plot for PCA and RDA analysis of *Populus trichocarpa* dataset.

**Figure S5:** PCA and scree plot of PCA on environmental variables of the *Populus trichocarpa* dataset.

